# Contemporary Enterovirus D68 strains show enhanced replication and translation at 37°C

**DOI:** 10.1101/2020.03.31.019380

**Authors:** Brendan D. Smith, Andrew Pekosz

## Abstract

Enterovirus D68 (EV-D68) emerged in 2014 as an important pathogen linked to severe lower respiratory disease and acute flaccid myelitis outbreaks. Historically associated with mild common-cold-like symptoms, clusters of severe disease attributed to EV-D68 appeared during a series of outbreaks in 2014, 2016, and 2018. Previous studies of historic EV-D68 strains demonstrated attenuated replication at temperatures of the lower respiratory tract (37°C), when compared to the upper respiratory tract (32°C). By testing a panel of historic and contemporary EV-D68 strains at 32°C and 37°C, we demonstrate that contemporary strains of EV-D68 undergo little to no attenuation at increased temperatures. Contemporary strains produced higher levels of viral proteins at 32°C and 37°C than historic strains, although both strains infected similar numbers of cells and had comparable amounts of replication complexes. IRES activity assays with dual-luciferase reporter plasmids demonstrated enhanced translation in recent EV-D68 strains mapped to regions of variability in the 5’ UTR found only in contemporary strains. Using an infectious clone system, we demonstrate that the translation advantage dictated by the 5’ UTR does not solely mediate temperature sensitivity. The strain-dependent effects of temperature on the EV-D68 life cycle gives insight into the susceptibility of the lower respiratory system to contemporary strains.

**IMPORTANCE:** Enterovirus-D68 (EV-D68) emerged in 2014 as a causative agent of biannual severe pediatric respiratory disease and acute flaccid myelitis (AFM). We show that recent EV-D68 viruses have gained the ability to replicate at 37°C. Enhanced virus protein translation seemed to correlate with enhanced virus replication at 37°C but other genetic factors are also contributing to this phenotype. An enhanced ability to replicate at core body temperature may have allowed EV-D68 to penetrate both lower in the respiratory tract and into the central nervous system, explaining the recent surge in severe disease associated with virus infection.

## INTRODUCTION

Human enterovirus D68 (EV-D68) has recently been associated with global outbreaks of severe acute respiratory illness and acute flaccid myelitis. This positive-sense, single-stranded RNA virus of the *Picornaviridae* family was first isolated in California from four pediatric patients in 1962. EV-D68 was found to be remarkably similar to human rhinovirus (HRV) in that it was acid labile, temperature sensitive, and transmitted from person-to-person via respiratory droplets [1-5]. The biological similarities between in EV-D68 and HRV led to its initial misclassification as human rhinovirus 87, which was corrected by genetic analysis and serum neutralization studies [6, 7]. Between 1970 and 2005, only 26 cases were reported to the CDC and the virus was generally associated with mild upper respiratory illness [8, 9].In 2014, unprecedented outbreaks of EV-D68 associated with severe respiratory disease and acute flaccid myelitis emerged throughout the world [3, 10-30]. Since 2014, there have been biannual patterns of severe disease associated with EV-D68 [31-50]. This unique pattern of recurring outbreaks and severe pathology sheds light on the importance of understanding evolutionary changes in contemporary EV-D68 strains.

The propensity of a respiratory pathogen to induce severe respiratory distress can be partly linked to its ability to replicate in the lower respiratory tract. The upper respiratory tract is approximately 32°C due to the intake of cool ambient air, while the lower airways are around 37°C, closer to core body temperature [51]. When respiratory viruses replicate poorly at 37°C, their infections are often restricted to the upper airways, limiting the potential for damage to the lower respiratory tract and ultimately mitigating disease severity. It is widely accepted that the reason most HRVs cause only mild upper respiratory illness is their preferential replication at cooler physiological temperatures [52-54].Temperature sensitivity and restriction to the upper airways has been harnessed for vaccine development and is the basis behind the cold-adapted live attenuated influenza vaccine [55, 56]. Historically, in vitro studies indicated EV-D68 replicates optimally at 33°C and is attenuated at higher physiological temperatures [1, 4, 5]. Given the recent outbreaks of lower respiratory disease associated with EV-D68, our lab sought to investigate EV-D68 temperature sensitivity in the context of historic and contemporary strains.

Picornaviruses have a positive-sense RNA genome approximately 7.5kb long that encodes a single open reading (ORF) that is translated into a single polypeptide. The genome also has 5’ and 3’ untranslated regions (UTRs) directly adjacent to the ORF. Both UTRs form secondary structure that are functionally involved in protein translation and genome replication. Picornaviruses utilize an internal ribosomal entry site (IRES) located in the highly structured 5’ untranslated region (UTR) of the viral genome for the initiation of viral protein translation [57, 58]. The IRES facilitates recruitment of host-cell translation machinery and initiation of translation in the absence of a 5’ methylated cap. Numerous studies have identified 5’ UTR mutations that impact IRES-mediated translation efficiency, which can be linked to temperature sensitivity, and neurotropism [57, 59, 60]. While some picornaviruses have been determined to have multiple open reading frames with an additional start codon in the 5’ UTR, this is not the case for EV-D68 [61, 62]. There has been substantial evolution in the EV-D68 5’ UTR since its initial discovery in 1962, most notably in the form of heightened variability and large deletions in the spacer region flanking the AUG start codon [63, 64]. Our studies identified a loss of temperature sensitivity in recent EV-D68 strains which was associated with increased translation of the viral genome. While the precise viral regions responsible for temperature sensitivity remain to be delineated, our results suggest the acquisition of efficient replication at 37°C preceded the emergence of EV-D68 as a global pathogen and may help explain its recent altered ability to induce respiratory and nervous system disease.

## MATERIALS AND METHODS

### Cell culture

Human Rhabdomyosarcoma (RD) cells were obtained from the American Type Culture Collection (ATCC) (CCL-136) and cultured in Gibco Minimal Essential Medium (MEM) supplemented with 10% fetal bovine serum (FBS), 100 U/mL penicillin, 100 μg/mL streptomycin, and 2 mM GlutaMAX (Gibco). The cells were grown at 37°C in a humidified environment supplemented to 5% CO2.

### Viruses

F02-3607 Corn (Corn-1963), US/MO/14-18947 (MO-2014), US/KY/14-18953, US/IL/14-18952 and AY426531 (Fermon-1962) were obtained from ATCC. USA/N0051U5/2012 (TN-2012) and US/MO/14-18949 (MO-2014/2) were provided by Dr. Suman Das of Vanderbilt University. Virus stocks were generated by inoculating a confluent T75 or T150 flask of RD cells at a multiplicity of infection (MOI) of 0.01 50% tissue culture infectious doses (TCID_50_) per cell for 1 hour at 32°C with rocking every 10-15 minutes. Virus inoculum was aspirated and replaced with infection media (IM-2.5% FBS, 100 U/mL penicillin, 100 μg/mL streptomycin, 2 mM GlutaMAX [Gibco]) followed by incubation at 32°C. After 24 hours of infection, initial IM was aspirated and replaced with fresh IM. When complete cytopathic effect (CPE) could be determined using a light microscope (3-5 days), the infected cells and supernatant were harvested, subjected to 2 freeze thaw cycles, and centrifuged at 600 x *g* for 10 minutes to remove all cell debris. The clarified supernatant was aliquoted and stored at -70C. Virus stock titers were determined using TCID_50_ and plaque assays described below.

### 50% Tissue culture infectious dose (TCID_50_) assay

RD cells were plated in 96-well plates, grown to full confluence, and washed with PBS+. Tenfold serial dilutions of the virus inoculum were made and 200 μL of dilution was added to each of 6 wells in the plate, followed by incubation at 32°C or 37°C for 5-7 days. Cells were then fixed by adding 100 μL of 4% formaldehyde in PBS per well and incubated at room temperature for at least 10 minutes before staining with Naphthol Blue Black solution. TCID_50_ calculations were determined by the Reed-Muench method [65].

### Plaque assays

Plaque assays were performed using 6-well plates of confluent RD cells. Serial 10-fold dilutions of the virus inoculum were generated in IM. Cells were washed twice with PBS containing calcium and magnesium (PBS+) and 200 μL of inoculum was added per well. The cells were incubated for 1 hour at 32°C with rocking every 10-15 minutes. The inoculum was removed and replaced with IM containing 1% agarose. Cells were then incubated at 32°C or 37°C for 5-7 days, fixed in 2% formaldehyde, and stained in a Naphthol Blue Black solution.

### Growth Curves

Multistep growth curves were performed on RD cells at an MOI of 0.001. Virus stocks were diluted to the appropriate concentrations in IM. Cells were inoculated with virus and incubated at 32°C with rocking every 10-15 minutes. Cells were then washed 2 times with PBS+ and incubated with fresh IM at 32 or 37°C. At the time points indicated, all supernatant was harvested and stored at -80°C. Infectious virus quantification at each time point was determined using TCID_50_ assays.

### Flow Cytometry

RD cells were washed with PBS+ and infected at an MOI of 5 for 1 hour at 32°C with rocking every 10-15 minutes. Infected cells were allowed to incubate for an additional 8 hours at 32°C or 37°C, at which time they were washed twice with PBS-, detached using trypsin, and fixed for 15 minutes at room temperature with 2% paraformaldehyde (PFA) in PBS. Cells were permeabilized with 0.2% Triton X-100 (Sigma) in PBS, blocked for 1 h in PBS with 3% normal goat serum and 0.5% bovine serum albumin (BSA; Sigma), and incubated for 1 h with anti-VP1 rabbit serum (1:200; GTX132313, Genetex) or dsRNA mAb (1:200; J2 anti-dsRNA IgG2a, Scicons), followed by incubation for 1 h with goat anti-rabbit AF647 (1:400; A-21224, Molecular Probes) or goat anti-mouse AF488 (1:200, A-11029, Molecular Probes). The cells were analyzed by flow cytometry (BD FACSCalibur) and data analyzed using FlowJo software.

### Phylogenetic Analysis and RNA secondary structure

Evolutionary trees were generated with all available EV-D68 complete genome sequences (408 sequences, December 11, 2017) from NCBI using entire genomes or 5’ UTRs only. Phylogenetic trees were generated using the Maximum-Likelihood tree analysis with 500 bootstrap replicates from MEGA-X software. Alignments shown were performed using Geneious 8.1.9 software.

Predicted RNA secondary structure for each domain was determined using both RNAstructure 6.1 and Geneious 8.1 software and referencing that of CBV [66]. The structures were visualized and edited using RNAstructure Structure Editor.

### IRES Activity Assay

Dual-luciferase plasmids were obtained from Genscript which used a CMV promoter to express and mRNA with the *renilla* luciferase gene, followed by the EV-D68 5’ UTR and firefly luciferase. RD cells were grown to 85-90% confluence in opaque 96-well plates overnight and plasmids were transfected into the cells for 4 hours at 37°C using TransIT-LT1 transfection reagent and incubated overnight at 32°C. The cells were then left at 32°C or moved to 37°C for 9 hours before read on a luminometer using the Dual-Luciferase Reporter Assay Step (Promega) according to manufacturer’s instructions. IRES activity was determined as the ratio of Firefly luciferase to *Renilla* luciferase.

### Infectious Clone System

Recombinant Corn-1963, MO-2014, and 5’ UTR chimeric viruses were generated using sequences available in GenBank. pUC57 vectors encoding a T7 transcription site followed by the full-length genome of interest, a 30-35 base poly-A tail, and *Cla*I restriction site for linearization were ordered from Genscript (Piscataway, NJ). Plasmids were fully linearized by incubation at 37°C for a minimum of 2 hours using *Cla*I restriction enzyme (New England Biolabs). Full-length genomic RNA was transcribed from the linearized plasmids using the MegaScript T7 Kit (Ambion). 4µg of transcribed RNA was mixed with DMRIE-C (Invitrogen) transfection reagent (1:2 µg:µL ratio) in Opti-MEM (Gibco), serial diluted in 4-fold increments (1µg to 0.016µg RNA final), and added to 85-90% confluent RD cells in 12-well plates for 2-4 hours at 37°C followed by an overlay of IM containing 1% agarose. The plates were incubated for up to 7 days with daily monitoring for plaques. Upon appearance, plaques were picked and plaque plugs placed in 500 µL of IM for use in generating seed stocks on RD cells.

### SDS-PAGE and western blotting

RD cells were infected at a high MOI for 1 hour at 32°C with rocking every 10-15 minutes and allowed to incubate for an additional 8 hours. Cells were washed twice with PBS and lysed using 1% SDS in PBS. Cell lysates were sonicated and clarified of any remaining cellular debris by centrifugation. Samples were mixed with 4X Laemmli buffer (Bio-Rad) containing dithiothreitol (DTT; ThermoFisher) and denatured at 100°C for 5 min. Samples were loaded alongside Precision Plus Protein Standards All Blue protein ladder (Biorad) into a 4-20% Mini-PROTEAN TGX Gel (Biorad) and run at 100 V for approximately 1 hour. Proteins were transferred to a PDVF membrane for 1 hour at 100V and blocked for 1 hour (5% blotting-grade blocker (Bio-Rad) in PBS containing 0.05% Tween 20). The membrane was stained for 1 h with mouse anti-VP3 (1:1000; GTX633706, Genetex) or mouse anti-β-actin (1:10000; Abcam), followed by staining for 1 h with anti-mouse AF647 (1:1000; A-28181, Molecular Probes). The membrane was washed 3 times for 5 minutes with PBS containing 0.05% Tween 20 with rocking between each step. Protein expression was analyzed using a FluorChem Q system. Cellular expression of viral proteins was normalized by β-actin expression.

### Statistical analyses

Replication kinetics of low MOI growth curves were analyzed by two-way ANOVA. TCID50 assays, FACS data, and IRES activity assays were analyzed using unpaired or multiple t-tests. All statistical analyses were performed in GraphPad Prism 8.0.1.

## RESULTS

### Effect of physiological temperature ranges on historic and contemporary EV-D68 strain infectious virus production

Historic strains of EV-D68 were previously shown to have optimal growth temperatures at 33°C and undergo attenuated growth at 37°C [1, 4, 5]. Given the recent rise in acute respiratory illness correlating with EV-D68, we tested a panel of historic and contemporary strains to determine if temperature sensitivity phenotypes differ between historic and contemporary strains. In agreement with the literature, historic strains of EV-D68 were attenuated at 37°C compared to 32°C (Fig. 1A), with TCID_50_ values reduced by 1000 to 100,000 fold. However, contemporary strains showed no to at most a 20 fold reduction in infectious virus titer when TCID_50_ assays are performed at 32°C or 37°C (Fig. 1A). When analyzing plaque assays incubated at 32°C or 37°C, contemporary strains showed more plaques of greater size at 37°C when compared to historic strains (Fig. 1B). In low MOI growth curves on RD cells (Fig. 1C), the Corn-1963 strain showed reduced infectious virus production at 37°C, while the MO-2014 produced equivalent infectious virus titers at both temperatures. At 32°C, MO-2014 produced higher infectious virus titers compared to Corn-1963, suggesting MO-2014 had improved replication at 32°C compared to Corn-1963. The data suggest contemporary strains of EV-D68 are better able to replicate at higher physiological temperatures than historic strains.

**Figure 1.**
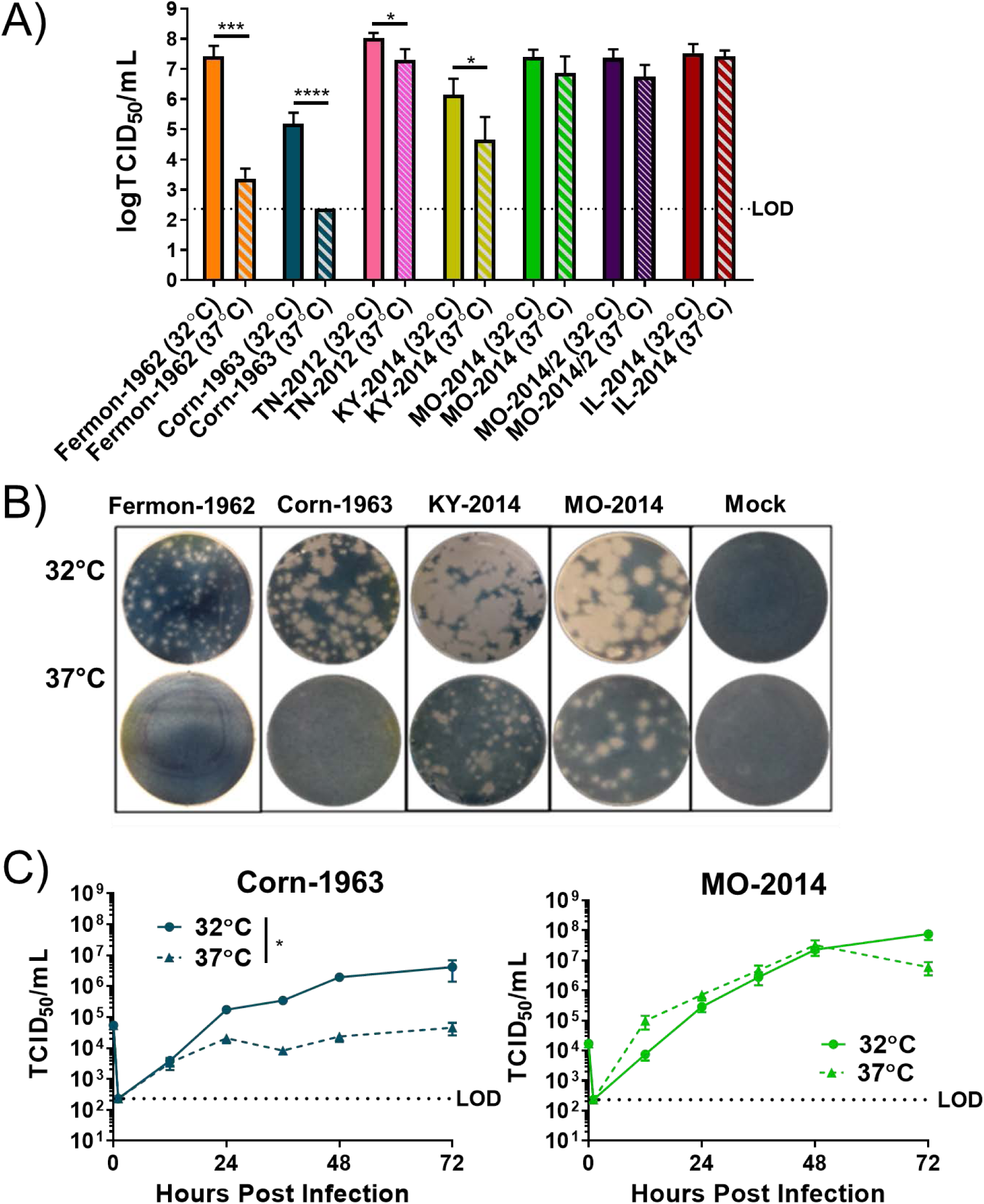
The effect of physiological temperature ranges on EV-D68 infectious virus production. (A) Infectious virus titers were determined with various EV-D68 virus strains at 32°C and 37°C using (A) TCID_50_ or (B) plaque assays. (C) RD cells were infected at an MOI of 0.01 and infectious virus titers (TCID_50_/mL) in the cell supernatants were determined at various times post infection. Statistical significance determined by unpaired t-test (TCID_50_) or 2-way ANOVA (growth curves) with *p<0.05, ***p<0.001, ****p<0.0001. The data shown are pooled from 3 experiments performed in triplicate for each experiment.

### Translation efficiency of contemporary and historic EV-D68 strains at physiological temperature ranges

To investigate the mechanisms controlling temperature restriction of EV-D68, we selected temperature sensitive Corn-1963 (historic) and temperature tolerant MO-2014 (contemporary) strains to delineate which stages of the EV-D68 life cycle were affected by temperature. After infecting RD cells (MOI of 5) for 9 hours at 32°C or 37°C, we detected viral RNA replication complexes using an antibody specific to dsRNA and viral protein using an antibody specific for EV-D68 viral capsid protein 1 (VP1). Using flow cytometry for analysis, both strains showed similar numbers of infected cells at both temperatures as detected by dsRNA positive cells (Fig. 2A) and nearly all of these infected cells were generating detectible viral protein (Fig. 2B). Mean fluorescence intensity was used to quantify relative amounts of dsRNA and VP1 on a per cell basis. The MFI of dsRNA was similar between strains (Fig. 2C). When RT-qPCR was used to quantify genome copies, there was a difference between Corn-1963 and MO02014 at both temperatures (Fig. 2E). Significant differences in VP1 expression were detected by flow cytometry, with the MO-2014 showing higher protein production at 32°C and 37°C compared to Corn-1963 (Fig. 2D). The differences in translation were further confirmed using a western blot assay and a monoclonal antibody to EV-D68 viral capsid protein 3 (VP3), which demonstrated increased VP3 at both 32°C and 37°C in MO-2014 infected cells (Fig. 2F). Based on the data, we inferred that while Corn-1963 and MO-2014 infected similar numbers of RD cells at either temperature, MO-2014 produced significantly more protein at both 32°C and 37°C.

**Figure 2.**
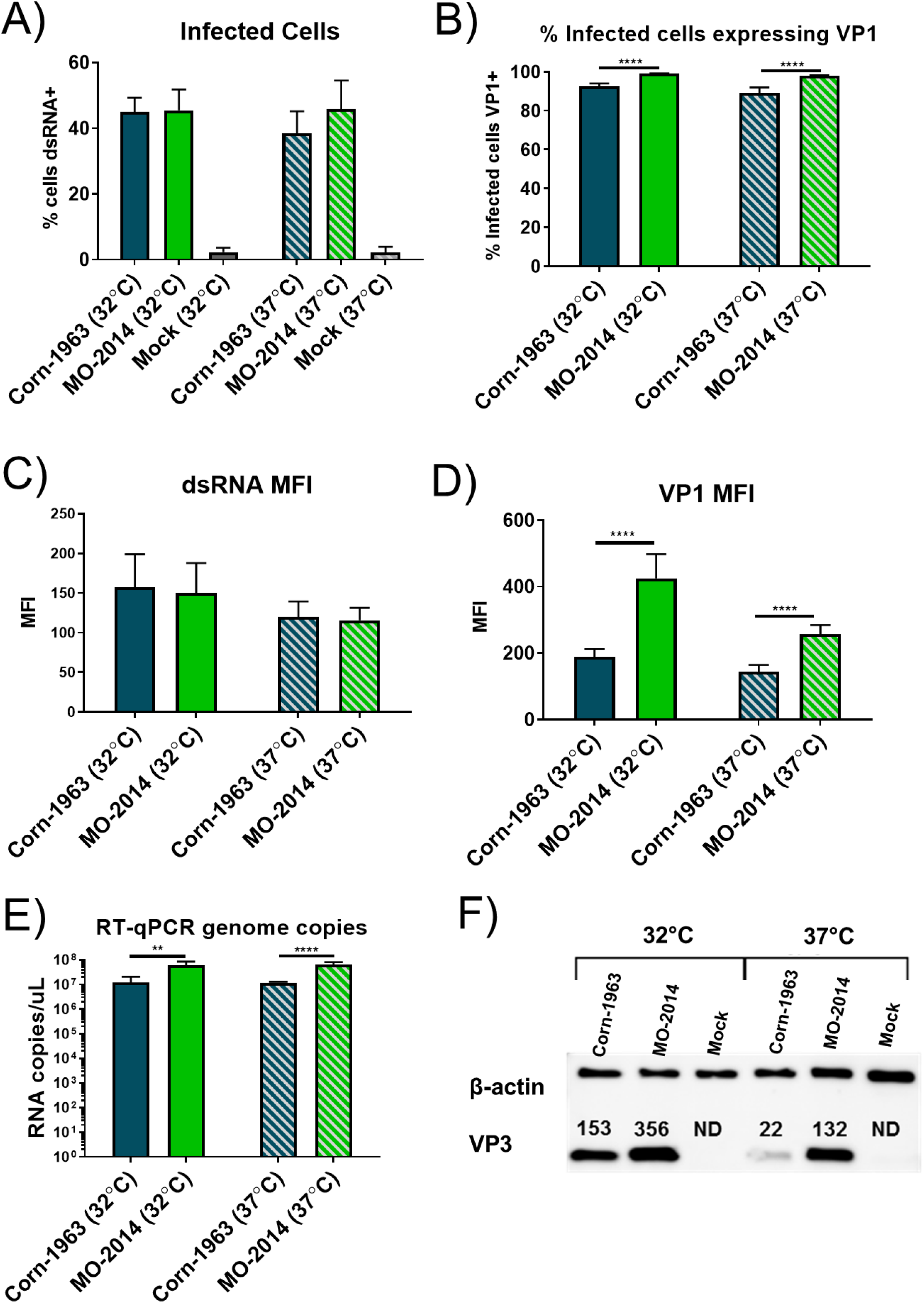
The effect of physiological temperature ranges on EV-D68 mRNA translation efficiency. RD cells were infected with Corn-1963 or MO-2014, or Mock (infection media) for 9 hours at an MOI of 5 and measured by flow cytometry for: (A) Percentage of infected cells (double-stranded RNA positive), (B) percentage of infected cells (gated to dsRNA+ cells) actively producing VP1 (VP1+), (C) relative amount of dsRNA in infected cells, measured by mean fluorescence intensity, and (D) relative amount of VP1 in infected cells. (E) EV-D68 genome copies were determined in virus infected cells by RT-qPCR. (F) Representative data of β-actin (loading control) and VP3 protein amounts analyzed by western blot in cell lysates generated at 9 hours post infection (MOI=5). VP3 band intensity was quantified using FluorChem Q and normalized to the intensity of the corresponding β-actin band (ND=not detected). Statistical significance determined by unpaired t-tests with *p<0.05, **p<0.01, ***p<0.001, ****p<0.0001. The data shown are pooled from 2-3 experiments performed in triplicate for each experiment.

### Assessment of IRES activity from strains ranging from 1963 to 2016

Since MO-2014 showed enhanced translation efficiency at 32°C and 37°C compared to the Corn-1963, we wanted to determine if IRES activity differed across a historical panel of virus strains. Sequences available on GenBank were selected that spanned all clades and a phylogenetic analysis was performed using either the IRES (Fig. 3A) or full genome (Fig. 3B) sequences. Representative 5’ UTR sequences from various EV-D68 strains were chosen for IRES activity analysis using a dual-luciferase plasmid encoding a CMV-promoter driven *renilla* luciferase ahead of the 5’ UTR of interest, followed by firefly luciferase. These plasmids were transfected into RD cells and IRES activity levels were determined at both 32°C and 37°C for each strain. At 32°C, all strains from 1999 or later had significantly higher levels of IRES activity compared to CORN-1963. At 37°C, five of the strains were significantly higher than Corn-1963, including three clade B strains from recent outbreaks, WY-2014, MO-2014, and MO-2016. The data indicate that the WY-2014, MO-2014 and MO-2016 5’ UTRs appear to have much greater ability to mediate translation at either 32°C or 37°C when compared to many historical EV-D68 strains.

**Figure 3.**
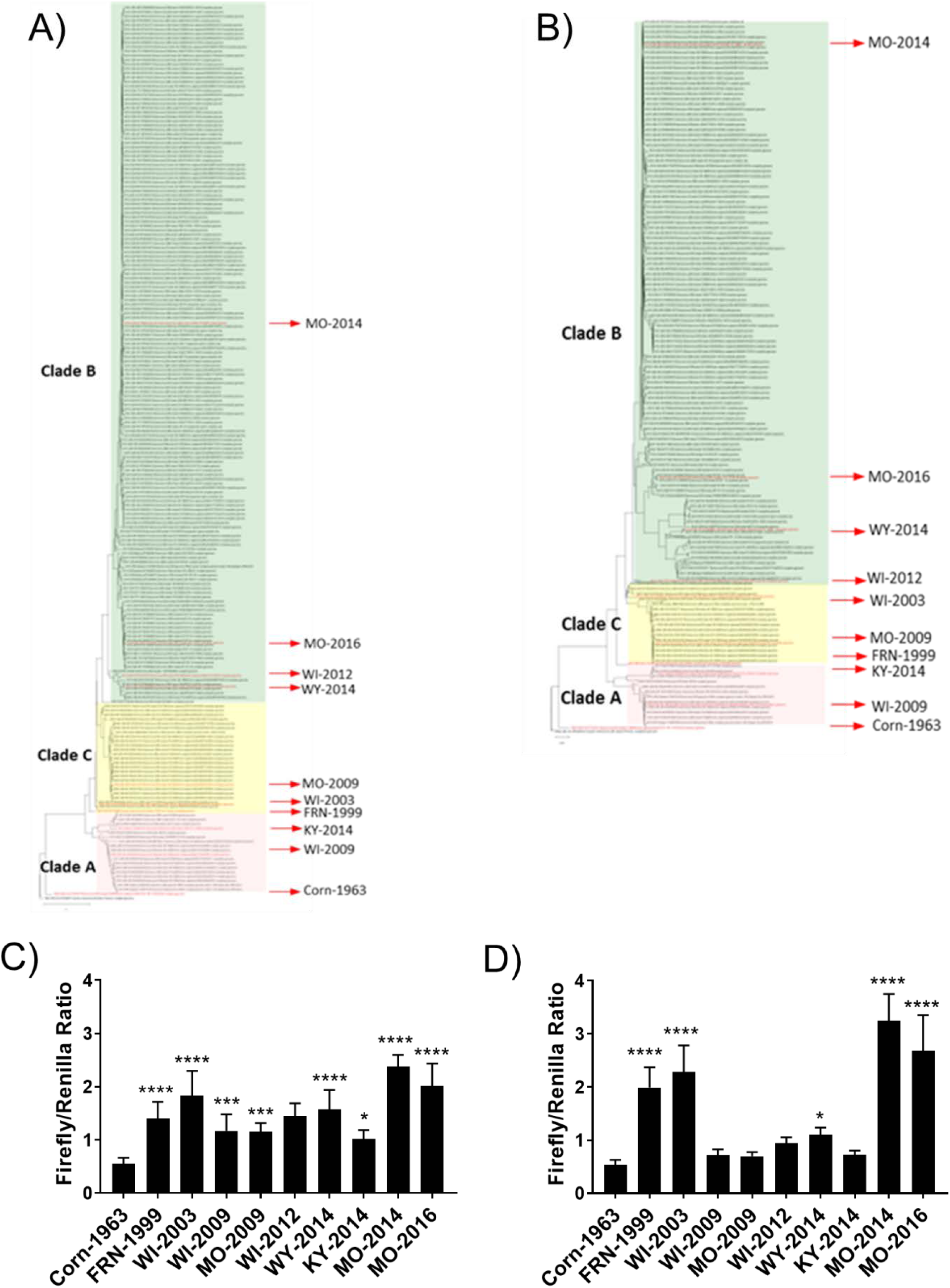
Changes in EV-D68 IRES activity from 1963 to 2016. Phylogenetic trees generated with all available EV-D68 complete genome sequences (408 sequences, December 11, 2017) from NCBI using (A) the entire genome or (B) 5’ UTR only. Trees were generated using the Maximum-Likelihood tree analysis with 500 bootstrap replicates from MEGA-X software. Various 5’ UTR sequences from historical EV-D68 sequences were selected (phylogeny indicated by red arrows on trees) and IRES activity analyzed at (C) 32°C or (D) 37°C using a dual-luciferase plasmid reporting system encoding the 5’ UTR EV-D68 sequences driving firefly luciferase activity (IRES mediated) compared to *renilla* luciferase activity (CMV-promotor mediated). Relative expression of firefly to *renilla* luciferase activity is graphed. Statistical significance determined by one-way ANOVA to Corn-1963 with *p<0.05, **p<0.01, ***p<0.001, ****p<0.0001. The data shown are pooled from 3 experiments performed in triplicate for each experiment.

### Genomic and structural comparison of Corn-1963 and MO-2014 and effects on IRES activity

We further analyzed the genomic and structural components of the 5’ UTR by aligning the corresponding sequences from Corn-1963 and MO-2014. The 5’ UTRs of Corn-1963 and MO-2014 share 83% homology with sporadic mutations that span the entire 5’ UTR. However, (Fig. 4A), there is a region of variation in bases 629-733 adjacent to the start codon, including a 35 base deletion previously reported to be in all contemporary EV-D68 strains. Of the 733 base 5’ UTR, 50% of the sequence variation is found in this variable region. The sequence variation between Corn-1963 and MO-2014 affect the predicted secondary structure (Fig. 4B) by inducing the formation of a stem loop structure (VII) in MO-2014 compared to Corn-1963. Using the dual-luciferase plasmid assay, we analyzed the contribution to IRES activity of both the 35 base deletion and the entire variable region of the 5’ UTR (Fig. 4C). Deleting the 35 bases in Corn-1963 increased IRES activity slightly. However, replacing the entire variable region with that of MO-2014 increased IRES activity, nearing levels of MO-2014. Adding in 35 bases to MO-2014 increased IRES activity slightly, but replacing the entire variable region with that of Corn-1963 resulted in a significant decrease in IRES activity. While the observed trends were detected at both 32°C and 37°C, the advantage of MO-2014 and its variable region penetrated more at 37°C. In fact, the ratio of IRES activity (MO-2014:Corn-1963) was significantly higher at 37°C, suggesting an advantage to MO-2014 at 37°C (Fig. 4D). Taken together, the enhanced IRES activity of MO-2014 is largely mediated by the 5’ UTR variable region which is more advantageous at 37°C.

**Figure 4.**
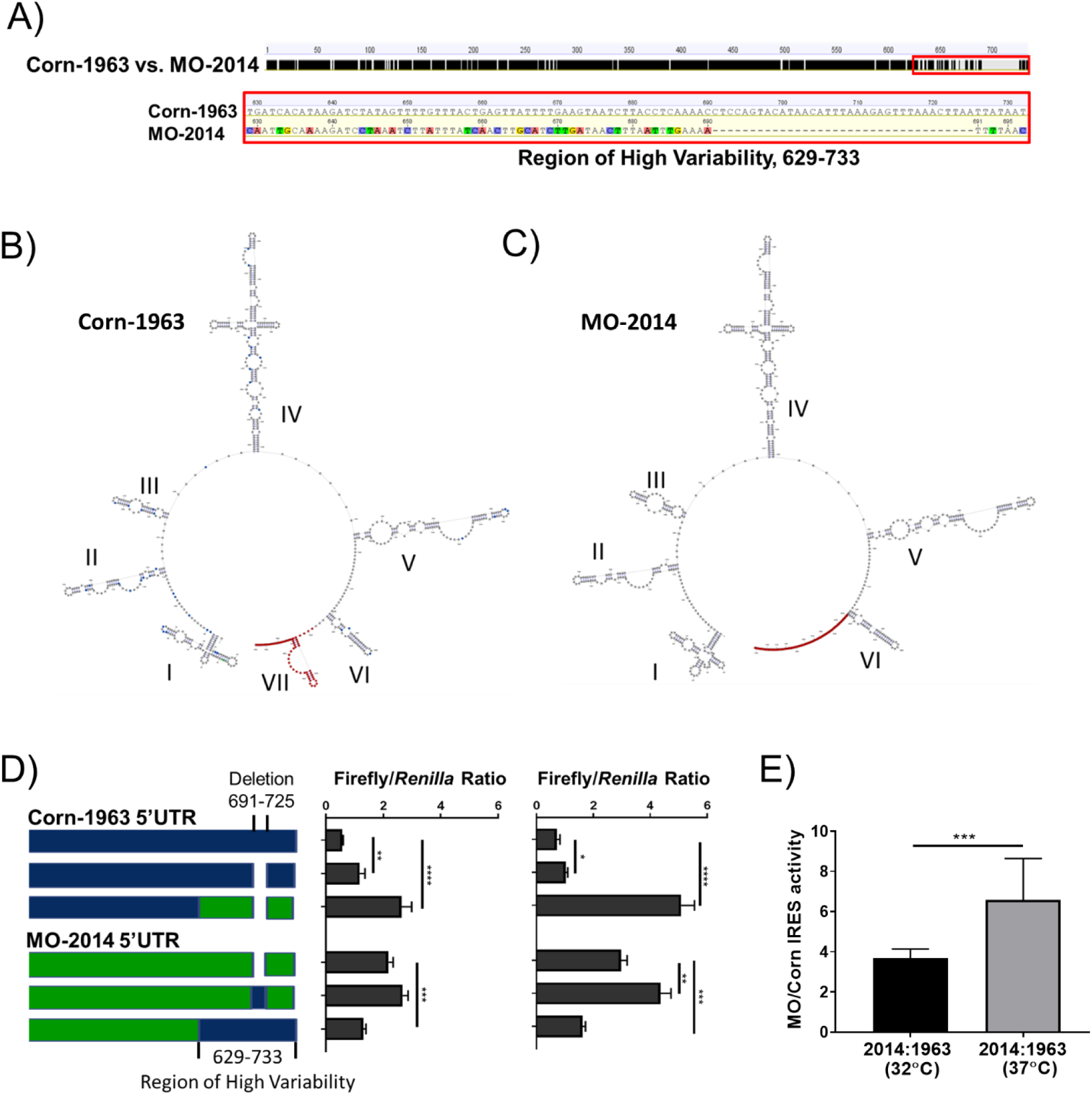
Comparison of Corn-1963 and MO-2014 IRES genetic features and effects on IRES activity. (A) Sequence alignment of Corn-1963 and MO-2014 5’ UTR (conserved sequence black, non-conserved in grey) identifies a 104 nt region of high variability (red box), including a 35 base deletion present in contemporary EV-D68 strains. (B) Predicted secondary structure of Corn-1963 and MO-2014 determined using RNAstructure 6.1 and Geneious 8.1.9 and visualized using RNAstructure StructureEditor software highlighting the region of high variability (red), point mutations (blue), and deletion (green) of MO-2014. (C) IRES activity measured using a dual-luciferase reporter system at 32°C (left) and 37°C (right) for Corn-1963 5’ UTR (blue), MO-2014 (green) and the chimeric 5’ UTRs. (D) Ratios of MO-2014 and Corn-1963 IRES activities were determined at 32°C (black) and 37°C (grey). Statistical significance determined by multiple or unpaired t-tests with *p<0.05, **p<0.01, ***p<0.001, ****p<0.0001. The data shown are pooled from 3 experiments performed in triplicate for each experiment.

### Replication of Corn-1963 and MO-2014 5’ UTR chimeric viruses at 32°C and 37°C

Because our IRES-activity assay indicated increased translation efficiency for MO-2014 that penetrated more strongly at 37°C, we generated chimeric recombinant viruses to test the influence of the 5’ UTRs on virus replication at 32°C and 37°C. TCID_50_ assays were performed at 32°C and 37°C on chimeric virus working stocks. The recombinant Corn-1963 (rCorn-1963) and MO-2014 (rMO-2014) viruses showed the same temperature sensitivity patterns as the natural isolates they were derived from in TCID50 assays (Fig. 5A). Surprisingly, replacing either the variable region or the entire 5’ UTR of either virus with that of the other had no effect on temperature sensitivity (Fig. 5A). Because TCID_50_ assays represent an endpoint assay, the effect of the 5’ UTR swaps were assessed in low MOI growth curves. Again, replacement of the variable region alone or the entire 5’ UTR had no effect on temperature dependent infectious virus production (Fig. 5B). Given that there was no change in the temperature sensitivity phenotypes involving infectious virus production, we determined the translation efficiency of the recombinant viruses using Western blotting for VP3. When the rCorn-1963 5’ UTR was replaced with that of rMO-2014, translation was increased (Fig. 5C; 32 to 57 at 32°C and 17 to 31 at 37°C) and the reciprocal swap resulted in a reduction of translation (Fig. 5C;117 to 42 at 32°C and 73 to 45 at 37°C). Replacing only the variable regions had a minimal impact on rCorn-1963, however, rMO-2014 lost translation efficiency when replaced with that of rCorn-1963. Taken together, the data demonstrate that that the 5’ UTR of MO-2014 increases viral protein translation in a recombinant virus when compared to the 5’ UTR of rCorn-1963. However, the 5’ UTR-mediated increase in translation efficiency is not the determining factor for EV-D68 temperature sensitive infectious virus production.

**Figure 5.**
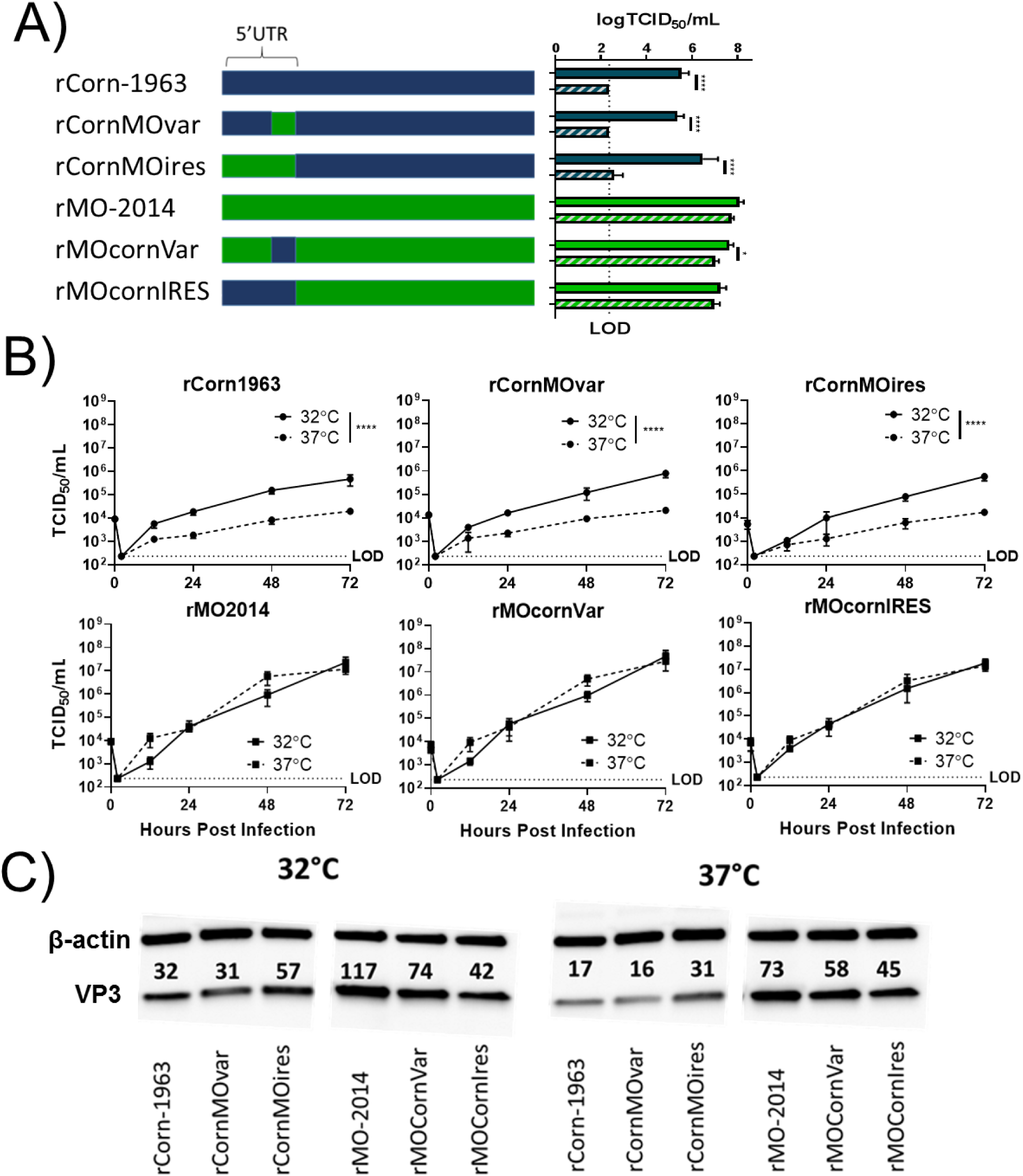
Effects of Corn-1963 and MO-2014 IRES chimeras on replication and viral protein translation. (A) Infectious virus titers were determined with Corn-1963 and MO-2014 chimeric IRES viruses at 32°C and 37°C using TCID_50_ assays. (B) RD cells were infected at an MOI of 0.01 and infectious virus titers (TCID_50_/mL) in the cell supernatants were determined at various time points post infection. (C) Representative data of β-actin (loading control) and VP3 protein amounts analyzed by western blot in cell lysates generated at 9 hours post infection (MOI=5). VP3 band intensity was quantified using FluorChem Q and normalized to the intensity of the corresponding β-actin band. Statistical significance determined by unpaired t-tests (TCID_50_) or 2-way ANOVA (growth curves) with *p<0.05, **p<0.01, ***p<0.001, ****p<0.0001. The data shown are for 2-3 experiments performed in triplicate for each experiment.

## DISCUSSION

EV-D68 has previously been shown to have an optimal growth temperature of 32°C compared to 37°C [1, 4, 5]. This temperature range is important because it represents the temperature of the human upper respiratory airways (32°C), which are cooled by inhalation of ambient air, and the lower respiratory airways (37°C), closer to core body temperature. Respiratory viruses incapable of replicating at higher physiological temperatures have correlated with mildly symptomatic infections of the upper respiratory tract [51]. HRV A and HRV B are two viruses with attenuated replication at higher physiological temperatures and are best known for causing common-cold symptoms [52, 53]. The recently discovered HRV C is not attenuated at 37°C and is associated with severe infections of the lower airways [67-71]. In EV-D68 outbreaks (2014, 2016, 2018), there have been correlations to increased acute respiratory illness (ARI) [21, 30, 72-77]. Our data indicate that contemporary strains of EV-D68 are better fit to replicate at 37°C than historic strains. Extrapolation of this data suggests that temperature is no barrier for the viruses to infect both upper and lower airways of the human respiratory tract.While recent studies have given insight into the ability of EV-D68 to infect neuronal cell lines in vitro and in animal models as it pertains to receptors, the mechanisms by which EV-D68 infects the central nervous system to cause AFM remain unclear [78-83]. Other picornaviruses capable of causing AFM such as poliovirus, EV-71, and CVB all replicate at high physiological temperatures. While temperature tolerance alone will not allow neuronal infections, our data suggests that contemporary strains of EV-D68 have overcome this barrier to efficient replication.

Picornaviruses utilize an internal ribosomal entry site (IRES) element located in the 5’ UTR of the virus genome for the initiation of viral protein translation. There have been various studies outlining the impacts of picornavirus 5’ UTR mutations on IRES-mediated translation efficiency. While some picornaviruses have been determined to have multiple start codons, this is not the case for EV-D68 [61, 62].There has been substantial evolution in the EV-D68 5’ UTR since its initial discovery in 1962. Comparing the genetic composition, there is 83% 5’ UTR sequence homology between our representative historic and contemporary strains and these differences do impact the overall predicted secondary structure (Fig. 4A and B). In addition to sporadic mutations throughout the 5’ UTR, there is a concentrated area of variability, including a 35 base deletion just prior to the translation start codon that is found in all contemporary EV-D68 strains. Due to the lack of sequences prior to the early 2000s, it is unknown when this sequence divergence took place. There is mounting evidence of EV-D68 recombination events that could potentially explain the heightened diversity in this specific region of the 5’ UTR [84]. Our dual-luciferase plasmid system gave the ability to test the impact of the 5’ UTR variability on translation by generating chimeric plasmids. Our data indicate that contemporary strains of EV-D68 have acquired increased translation efficiency driven by evolution in the 5’ UTR. The 35 base deletion did not have much of an effect on the historical strains translation efficiency, however, incorporation of the entire region of high variability increased translation significantly (Fig. 4C). Predictive secondary modeling suggest that the variable region of the contemporary strains lead to an additional stem-loop after domain VI (Fig. 4A). The effects of additional secondary structure in this region remains unknown, but could play a role in recruitment of translation initiation factors.

While our data suggest that contemporary strains of EV-D68 have a translation efficiency advantage over historic strains based on our bicistronic reporter assays, chimeric viruses were necessary to determine the impact it may have on infectious virus production and replication kinetics. Swapping the variable region or the entire 5’ UTRs of Corn-1963 and MO-2014 did not phenotypically change their temperature sensitivity. That is, Corn-1963 encompassing components of MO-2014 remained attenuated at 37°C, while MO-2014 with the 5’ UTR of Corn-1963 remained temperature tolerant (Fig. 5A and 5B). We confirmed that the 5’ UTR swaps do alter translation efficiency (Fig. 5C) in the context of a virus infection but the lack of an effect of 5’UTR swaps on the temperature sensitivity of infectious virus production indicates other viral sequences contribute to temperature tolerance (Fig. 5D). There are many other benefits that may arise by increased translation. Picornaviruses, including EV-D68, have been show to utilize their encoded proteases to manipulate the host-cell immune function [85, 86]. Increased translation efficiency and the generation of more of these proteases could allow for a more robust immune evasion. Increased translation efficiency may also play a role in disease pathogenesis.

Our results suggest that determinants outside the 5’ UTR control temperature depending infectious virus production. Recent literature suggests that EV-D68 temperature sensitivity may be linked to the viral capsid protein VP1 [4]. Specifically, by taking a temperature tolerant EV-D94 virus and swapping the VP1 with that of the prototype EV-D68 strain (Fermon-1962), the virus becomes temperature sensitive. While interesting, the reverse swap (EV-D94 VP1 into EV-D68) was not able to be rescued, and therefore a temperature sensitivity phenotype swap for EV-D68 has not yet been successful. Generating a full panel of genomic swaps between the representative strains used in our studies investigating the role of structural and non-structural genes would lend insight into their role in temperature tolerance. Additionally, while the 5’ UTR contains the IRES element, the 3’UTR for picornaviruses also contains secondary structure and has been shown to influence translation and replication [87-90]. While extensive studies have not been done previously for EV-D68 3’UTR, one can postulate that similar interactions may occur between its 5’ and 3’ UTRs. Including an assessment of the 3’UTR in chimeric viruses may lend insight into its involvement in temperature sensitive infectious virus production.

The data from these studies provide a functional basis for the genetic evolution that has been reported for the 5’ UTR of contemporary EV-D68 strains. Additionally, all contemporary strains of EV-D68 tolerate replication at 37°C better than historic strains. This provides evidence that contemporary strains may have increased their ability to infect the lower respiratory tract and CNS – sites of the body normally at 37°C. Continued research into the mechanism of contemporary strain temperature tolerance is essential for a full understanding of recent severe EV-D68 outbreaks. Given the continued biannual patterns associated with severe disease since 2014, understanding all potential contributors is essential.

## ACKNOWLEDGEMENTS

We would like to thank Dr. Suman Das at Vanderbilt University for kindly providing TN-2012 and MO-2014/2 strains. Thank you to Dr. Katherine Fenstermacher and Dr. Elizabeth Troisi for their initial work with our EV-D68 virus stocks and Alysha Ellison for her help with generating rMO-2014. We thank Harrison Powell of the Pekosz laboratory for his support with flow cytometry. We would like to thank members of the Pekosz laboratory, Dr. Sabra Klein and members of the Klein laboratory, Dr. Kimberly Davis and members of the Davis laboratory, and Dr. William Jackson and members of the Jackson laboratory for their critical discussions of the data. This work was supported by HHSN272201400007C (AP) and T32 AI007417 (BS).

